# Identification of microRNAs in the Lyme disease vector *Ixodes scapularis*

**DOI:** 10.1101/2022.04.14.488330

**Authors:** Deepak Kumar, Latoyia P. Downs, Monica Embers, Alex Flynt, Shahid Karim

## Abstract

MicroRNAs (miRNAs) are a class of small non-coding RNAs involved in many biological processes, including the immune pathways that control bacterial, parasitic, and viral infections. Pathogens probably modify host miRNAs to facilitate successful infection, so they might be useful targets for vaccination strategies. There are little data on differentially expressed miRNAs in the black-legged tick *Ixodes scapularis* after infection with *Borrelia burgdorferi*, the causative agent of Lyme disease in the United States. Small RNA sequencing and qRT-PCR analysis were used to identify and validate differentially-expressed *I. scapularis* salivary miRNAs. Small RNA-seq yielded 133,465,828 (≥18 nucleotides) and 163,852,135 (≥18 nucleotides) small RNA reads from *Borrelia*- infected and uninfected salivary glands for downstream analysis using the miRDeep2 algorithm. Two hundred and fifty-four miRNAs were identified across all datasets, 25 of which were high confidence and 51 low confidence known miRNAs. Twenty-three miRNAs were differentially expressed in uninfected and infected salivary glands: 11 were upregulated and 12 were downregulated upon pathogen infection. This study provides new insights into the miRNAs expressed in *I. scapularis* salivary glands and pave the way for their functional manipulation to prevent or treat *B. burgdorferi* infection.

## Introduction

Small non-coding RNAs (sncRNAs) regulate genes at the post-transcriptional level in animals, plants, and arthropods including ticks [1-5]. MicroRNAs (miRNAs) are a class of sncRNA between 18 and 25 nucleotides in length, and they are now known to be important in arthropod immunity and host-pathogen interactions through their involvement in several cellular processes including development, immunity, and pathogen responses in arthropods [2, 6-10]. In animals, miRNAs regulate post-transcriptional gene expression, most often by binding to the 3’-untranslated region (3’-UTR) of target genes. While perfect complementarity of 2-8 nucleotides at the 5’ end of the miRNA (seed region) is necessary for miRNA regulation, the remaining sequence can harbor mismatches or bulges [2, 11, 12]. miRNAs are transcribed as primary miRNA transcripts before processing by Drosha and Pasha proteins into a pre-miRNAs. These are then exported to the cytoplasm and processed by Dicer into mature miRNAs, which are loaded onto the microRNA-induced silencing complex (miRISC) before targeting complementary mRNA for degradation [13, 14].

Several proteomics and transcriptomics studies have identified differentially expressed transcripts and proteins in uninfected and infected ticks [1, 15-18], but few have investigated the role of pathogens in the differential modulation of the post-transcriptional tick and host machinery. Although there are over 800 tick species and despite their importance as vectors of human and animal diseases, ticks are underrepresented in available miRNA resources. For example, the miRbase database contains 49 *Ixodes scapularis* miRNAs and 24 *Rhipicephalus microplus* miRNAs, while MirGeneDB 2.1 contains 64 *Ixodes scapularis* miRNAs. However, smallRNA sequencing (RNA-seq) and computational approaches are accelerating the discovery of miRNAs from species with incomplete genome sequencing, assembly, and annotation.

*Ixodes scapularis* is a primary vector of human pathogens including the Lyme disease agent *Borrelia burgdorferi*, which infects vertebrates and ticks through evolved complex mechanisms. There is now a good understanding of tick immune pathways and their interactions with *B. burgdorferi* [19-21], but it is still uncertain how *B. burgdorferi* avoids clearance. *B. burgdorferi* must traverse tick salivary glands during transmission [71]. Since saliva/salivary gland proteins can enhance *B. burgdorferi* transmission into the vertebrate host, characterization of molecular interactions at the tick bite site and the tick salivary glands is expected to facilitate vaccine development [22, 23], since promoting immunity against tick salivary proteins could neutralize tick bites and pathogen transmission. Once *B. burgdorferi* is acquired by ticks from infected hosts, it resides in the tick gut and only migrates to the salivary gland during subsequent blood feeding, which generally lasts for 3-7 days [72, 73, 74, 75]. While it is known that other tick-borne pathogens such as *Anaplasma marginale* must replicate inside salivary glands for efficient transmission [76], the details of *B. burgdorferi* replication are less well understood. Indeed, borrelial spirochetes invade the tick salivary gland via an unknown mechanism [71] and might be carried to the host dermis via tick saliva. Several salivary gland genes are upregulated in *B. burgdorferi*-infected *Ixodes scapularis* nymphs compared with uninfected ones [24], suggesting a significant role for salivary gland gene regulation in *B. burgdorferi* infection and transmission.

To fill the knowledge gap on miRNA expression in *Ixodes scapularis* salivary glands, here we performed miRNA profiling of partially fed *B. burgdorferi*-infected and uninfected tick salivary glands to identify miRNAs that might play a role in *B. burgdorferi* survival, colonization, transmission, and host immunomodulation. In doing so, we detected 254 miRNAs, of which 25 were high confidence miRNAs and 51 low confidence miRNAs. Forty-one of the identified miRNAs were present as *I. scapularis* miRNAs in miRBase (v22.1). Gene ontology and network analysis of target genes of differentially expressed miRNAs predicted roles in metabolic, cellular, development, cellular component biogenesis, and biological regulation processes. Several KEGG pathways including sphingolipid metabolism; valine, leucine and isoleucine degradation; lipid transport and metabolism; exosome biogenesis and secretion; and phosphate-containing compound metabolic processes were predicted as targets of differentially expressed miRNAs.

## Materials and Methods

### Ethics statement

All animal experiments were performed in strict accordance with the recommendations in the NIH Guide for the Care and Use of Laboratory Animals. The Institutional Animal Care and Use Committee of the University of Southern Mississippi approved the protocol for blood feeding of field-collected ticks (protocol # 15101501.1).

### Ticks and tissue dissections

Ticks were purchased from the Oklahoma State University Tick Rearing Facility. Adult male and female *I. scapularis* were kept according to standard practices [25] and maintained in the laboratory as described in our previously published work [26, 27]. Unfed female adult *I. scapularis* were infected with laboratory grown *B. burgdorferi* strain B31.5A19 using the capillary feeding method at Tulane National Primate Research Center [28]. *Borrelia* infected and uninfected ticks (n=45 in each group) were placed on each ear of a rabbit host for tick blood-feeding. Blood-fed adult female *I. scapularis* were dissected within 60 min of removal from the rabbit. Tick tissues were dissected and washed in M-199 buffer as described previously [77]. Salivary glands and midguts from individual *I. scapularis* were stored in RNAlater (Life Technologies, Carlsbad, CA, USA) at −80°C until use.

### RNA isolation, cDNA synthesis, and PCR-based *B. burgdorferi* detection in tick tissues

The TRIzol method was used for RNA extraction from individually-dissected midgut tissues, and cDNA was synthesized as described previously [29, 78]. *B. burgdorferi* was detected in tick midguts using the *flaB* gene in a PCR assay [29, 30]. After testing for *B. burgdorferi* infection in tick midguts, the corresponding salivary gland tissues from the same uninfected/infected ticks (n=10 salivary glands from each group) were pooled in separate tubes and RNA isolated using the TRIzol method [78].

### Small RNA sequencing (RNA-seq)

Small RNA libraries were made using the Illumina TruSeq Kit following the manufacturer’s protocol (Illumina, San Diego, CA). Briefly, short adapter oligonucleotides were ligated to each end of the small RNAs in the sample, cDNA made with reverse transcriptase, and PCR used to add sample-specific barcodes and Illumina sequencing adapters. The final concentration of all sequencing libraries was determined using a Qubit fluorometric assay (Thermo Fisher Scientific, Waltham, MA), and the DNA fragment size of each library was assessed using a DNA 1000 high-sensitivity chip on an Agilent 2100 Bioanalyzer (Agilent Technologies, Santa Clara, CA). After purification by polyacrylamide gel electrophoresis, sample libraries were pooled and sequenced on an Illumina NextSeq 500 (single end 36 base) using the TruSeq SBS kit v3 (Illumina) and protocols defined by the manufacturer. Four small RNA libraries of clean and *B. burgdorferi*- infected, partially fed, and pooled salivary glands were sequenced. RNA library preparation and indexing were performed at the UMMC sequencing facility.

### Data analysis

A schematic of the experimental plan and data analysis is shown in **Supplementary Figure S1**. miRDeep2 v.2.0.0.8 [31, 32], was used to process RNA-seq data. To predict novel miRNAs, the reads from all samples were combined. The mapper function of miRDeep2 first trims the adapter sequences from the reads and converts the read files from FASTQ to FASTA format. Reads shorter than 18 bases were discarded and the remaining reads mapped to the *I. scapularis* reference genome using default miRDeep2 mapper function parameters. Reads mapping to the genome were used to predict novel miRNAs. The *Drosophila melanogaster* genome was also used as another reference genome, and mapped reads were aligned to available miRNAs of *D. melanogaster* in miRBase (v22) and quantified. Reads were mapped to the reference genomes of *D. melanogaster* and *I. scapularis* and locations of potential miRNA read accumulations identified. The regions immediately surrounding the mapped reads were examined for miRNA biogenesis features including mature miRNAs, star and precursor reads, and stem-loop folding properties. miRDeep2 models the miRNA biogenesis pathway and uses a probabilistic algorithm to score compatibility of the position and frequency of sequencing reads with the secondary structure of the miRNA precursor.

For miRNA expression, a count table was generated using bedtools multicov, which counts alignments from indexed BAM files that overlap intervals in BED files provided from the miRDeep2 analysis.

### ISE6 cell culture, infection with *B. burgdorferi*, and validation of differentially expressed miRNAs by qRT-PCR

Differentially expressed miRNAs in RNA-seq data were validated by qRT-PCR in the ISE6 cell line derived from *I. scapularis* embryos cultured and maintained as suggested by Munderloh and Kurtti [33]. The *B. burgdorferi* (strain B31) isolate was a kind donation from Dr. Monica Embers (Tulane University, Covington, LA). Cultures were grown and maintained as suggested previously [28]. Once ISE6 cells reached confluency, they were infected with *B. burgdorferi* as described previously [34]. Briefly, ISE6 cells were inoculated with the supernatant of a log phase *B. burgdorferi* culture at a multiplicity of infection of 50 and incubated for 24 hours before harvesting. RNA was isolated using the TRIzol method and the expression of miRNAs analyzed by qRT-PCR with the Mir-X miRNA qRT-PCR TB Green kit (Takara Bio Inc, Kusatsu, Shiga, Japan; catalog #638316). qRT-PCR conditions were an initial denaturation at 95°C for 10 mins then 40 cycles of 95°C for 5 secs, 60°C for 20 secs.

### Normalization, differential expression (DE), and statistical analysis of miRNAs between uninfected and infected salivary glands

Differential expression (DE) analysis of identified miRNAs was performed using the interactive web interface DeApp [35]. Low expression genetic features were removed after alignment if the counts per million (CPM) value was ≤1 in less than two samples. Sample normalization and a multidimensional scaling (MDS) plot are shown in **Supplementary Fig. S2**. DE analysis was performed with edgeR with a false discovery rate (FDR) adjusted *p*-value of 0.05 and minimum fold-change of 1.5. DeApp displays a dispersion plot showing the overall DE analysis along with statistical significance (*p*-value, FDR adjusted *p*-value) and a volcano plot corresponding to the specified parameters and cutoff values.

### *In silico* mapping of *B. burgdorferi*-infected small RNA sequences to the *Borrelia burgdorferi* genome

*In silico* mapping of small RNA sequences of *B. burgdorferi*-infected salivary glands to the *Borrelia burgdorferi* genome (GCF_000181575.2_ASM18157v2_genomic.fna) detected 18,165 *Borrelia burgdorferi* sequences, but no miRNAs of *B. burgdorferi* origin were predicted by miRDeep2 analysis.

### Prediction of target genes, proteome re-annotation, gene ontology (GO), and KEGG enrichment analyses

TargetSpy [36], MIRANDA [37], and PITA [38] were used in miRNAconsTarget from sRNAtoolbox to predict genes regulated by tick salivary gland miRNAs up- or downregulated in the *in silico* analysis [39]. Targets common to all three programs were further considered. *In silico* target prediction provided a high number of false positives, but cross-species comparisons and combinatorial effects reduced this number [40]. Target gene networks and KEGG pathways significantly enriched for target genes were extracted using the STRING [41] output. PANNZER2[42] was used to functionally re-annotate the proteome of up- or downregulated genes (targets of detected miRNAs), and WEGO [43] was used to analyze and plot gene ontology (GO) annotations.

## Results and Discussion

### Profile characteristics of small RNA libraries

There were 216,292,174 raw small RNA reads from *B. burgdorferi*-infected salivary glands and 212,542,697 from uninfected salivary glands. After adapter trimming and removal of short reads (≤18 nucleotides (nt)), 133,465,828 small RNA reads were available from *B. burgdorferi*-infected samples and 163,852,135 from uninfected samples for downstream analysis. The read length distribution shows the types of small RNAs present in *B. burgdorferi*-infected and uninfected salivary gland samples. Two main peaks at 22 nt (miRNAs/siRNAs) at 29 nt (piRNAs) were distinguishable in *B. burgdorferi*-infected and uninfected samples (**Figure 1A**). There were ∼5 × 10^7^ and ∼2.5 × 10^7^ 22 nt miRNA sequences in uninfected and infected salivary gland samples, respectively. The read length distribution in uninfected samples was comparable to infected samples.

**Figure 1.**
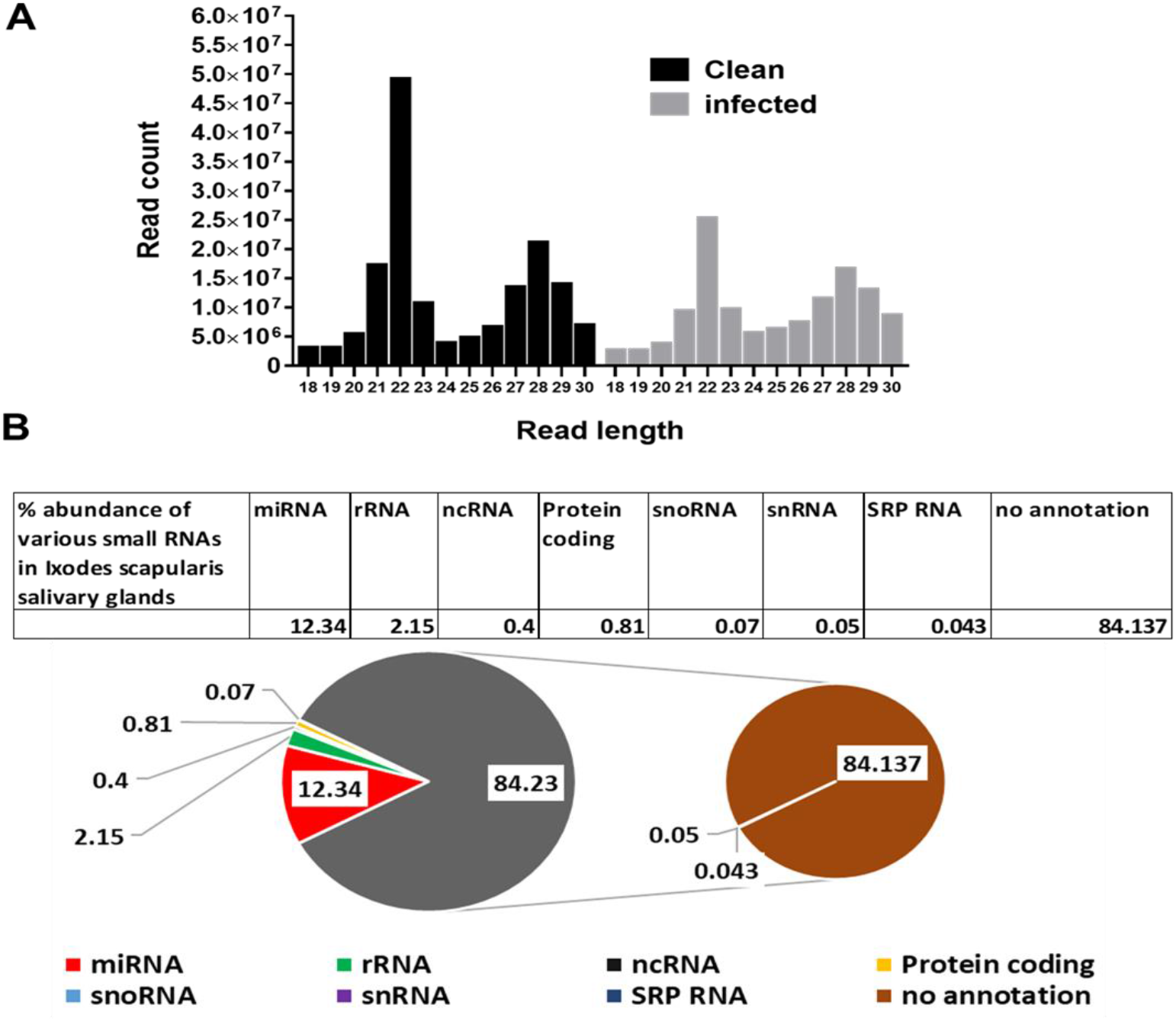
(A) Small RNA sequence length distribution in uninfected (clean) and *Borrelia burgdorferi*-infected *Ixodes scapularis* salivary glands. MicroRNAs are twenty-four (22) nucleotides in length. Clean, uninfected salivary glands; infected, *Borrelia burgdorferi*-infected salivary glands. (B) Summary of the abundance of combined small RNA reads from *Ixodes scapularis* salivary glands matching various small RNA categories. Reads from uninfected and *Borrelia burgdorferi* infected samples are combined together for abundance calculation.

### Other small RNA categories

A summary of reads matching various small RNAs categories from *B. burgdorferi*-infected and uninfected salivary glands are shown in **Figure 1B**. Other small RNAs include signal recognition particle rRNAs, ncRNAs, protein-coding RNAs, snoRNAs, snRNA SRP RNAs, and not annotated small RNAs. Of the total small RNA reads from *Ixodes scapularis* salivary glands, 12.3% were miRNAs, 2.2% rRNAs, 0.4% ncRNAs, 0.8% protein coding, 0.07% snoRNAs, 0.05% snRNAs, 0.04% SRP RNAs, and 84.3% reads were not annotated.

### MicroRNA profiling of infected and uninfected salivary glands and identification of novel tick miRNAs

**Fig. 2A** shows the basic hairpin-loop structure of an miRNA and other parameters (Dicer cut overhangs, total read count, mature read count, loop read count, total read count, randfold score, and total score) used to determine whether a hairpin-loop structured RNA is an miRNA. Using miRDeep2, 254 miRNAs were predicted in the salivary gland libraries. By adopting a conservative approach, miRNAs were categorized as high (n=25) and low confidence (n=51) miRNAs (**Fig. 2B**) based on standard criteria [79]. Of 49 *Ixodes scapularis* miRNAs present in miRBase, 41 were detected in our data. Of the 254 *Ixodes scapularis* miRNAs, several had homologs present in *D. melanogaster*. The details of the identified miRNAs and their miRDeep2 scores are presented in **Supplementary Table S1**.

**Figure 2.**
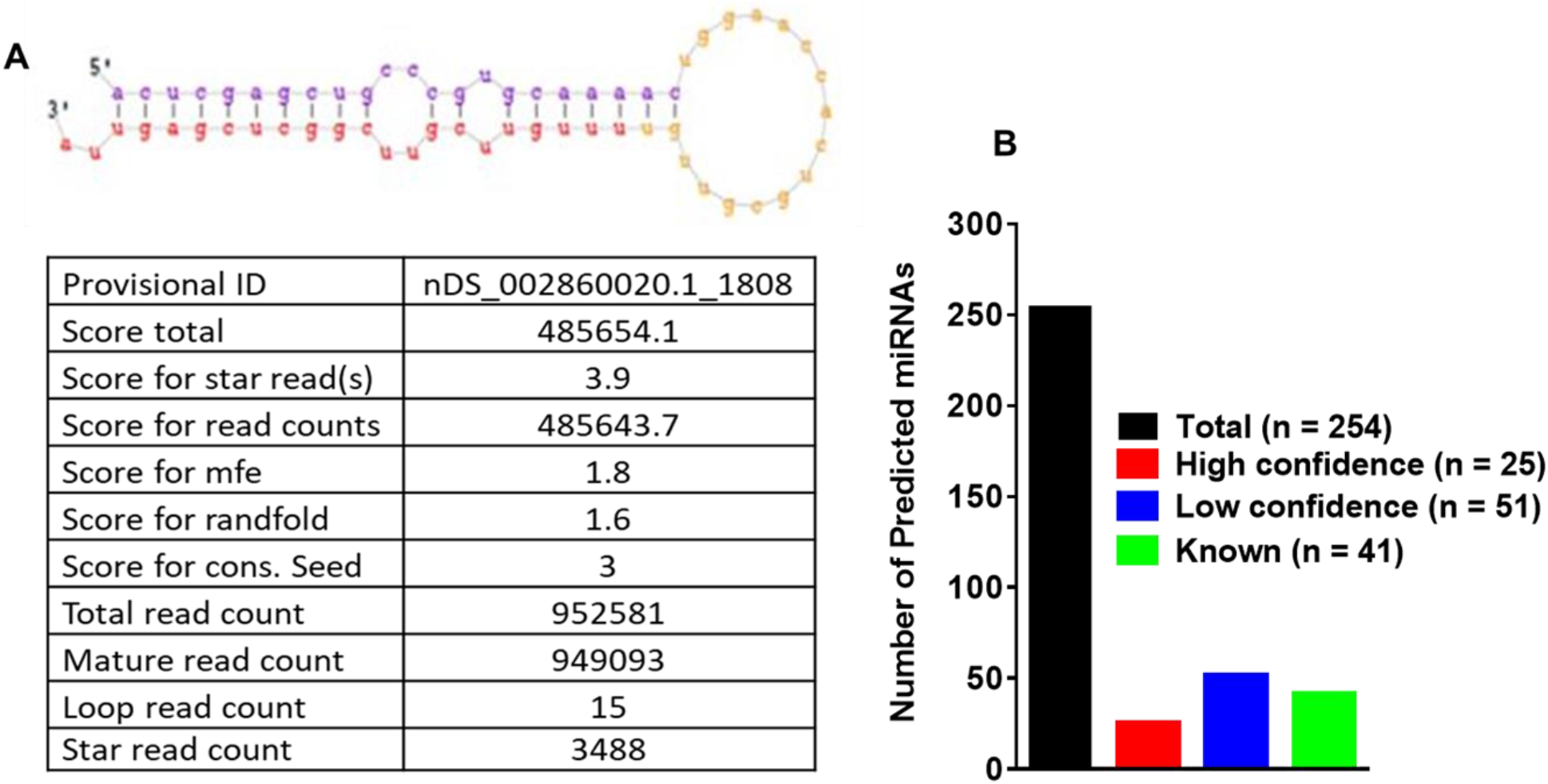
(A) Basic stem-loop structures of predicted microRNAs. miRDeep2 was used to identify potential miRNA precursors based on nucleotide length, star sequence, stem-loop folding, and homology to the *Ixodes scapularis* reference genome. Shown are the predicted stem-loop structures (yellow), star (pink), and mature sequences of predicted miRNAs (red) in the salivary glands of *Ixodes scapularis* ticks. (B) Annotation of predicted miRNAs in partially fed uninfected (clean) and *Borrelia burgdorferi*-infected *Ixodes scapularis* salivary glands. 254 microRNAs were predicted in infected and uninfected salivary gland tissues. 25 were categorized as high confidence (n = 25) and 51 as low confidence (n=51) based on standard criteria and a conservative approach. Out of 49 *Ixodes scapularis* miRNAs available in miRbase (v22.1), 41 were detected in this study.

### *In silico* DE analysis of miRNAs in *B. burgdorferi*-infected salivary glands

In DE analysis, 11 miRNAs were upregulated and 12 were downregulated in infected salivary glands compared with uninfected salivary glands (**Figure 3A, B; Table 1**). Several of the identified tick miRNAs were conserved in *D. melanogaster* (dme-miR-375-3p, dme-miR-993-5p, dme-miR-12-5p, dme-bantam-3p, dme-miR-100-5p, dme-miR-8-3p, and dme-miR-304-5p). miRNAs downregulated in *B. burgdorferi*-infected salivary glands relative to uninfected salivary glands were isc-miR-153, isc-miR-1 and isc-miR-79, nDS_002871802.1_376, nDS_002633080.1_31953, nDS_002549652.1_43448, nDS_002537755.1_45711, nDS_002537755.1_45739, nDS_002861213.1_1750, nDS_002763926.1_14582, and nDS_002548557.1_43749, while upregulated miRNAs were isc-miR-317, isc-miR5310, isc-miR- 2001, isc-miR-5307, isc-miR-71, isc-miR-87, nDS_002784743.1_12165, nDS_002664372.1_28027, nDS_002716687.1_21278, nDS_002620414.1_33611, and nDS_002680650.1_26098.

**Figure 3.**
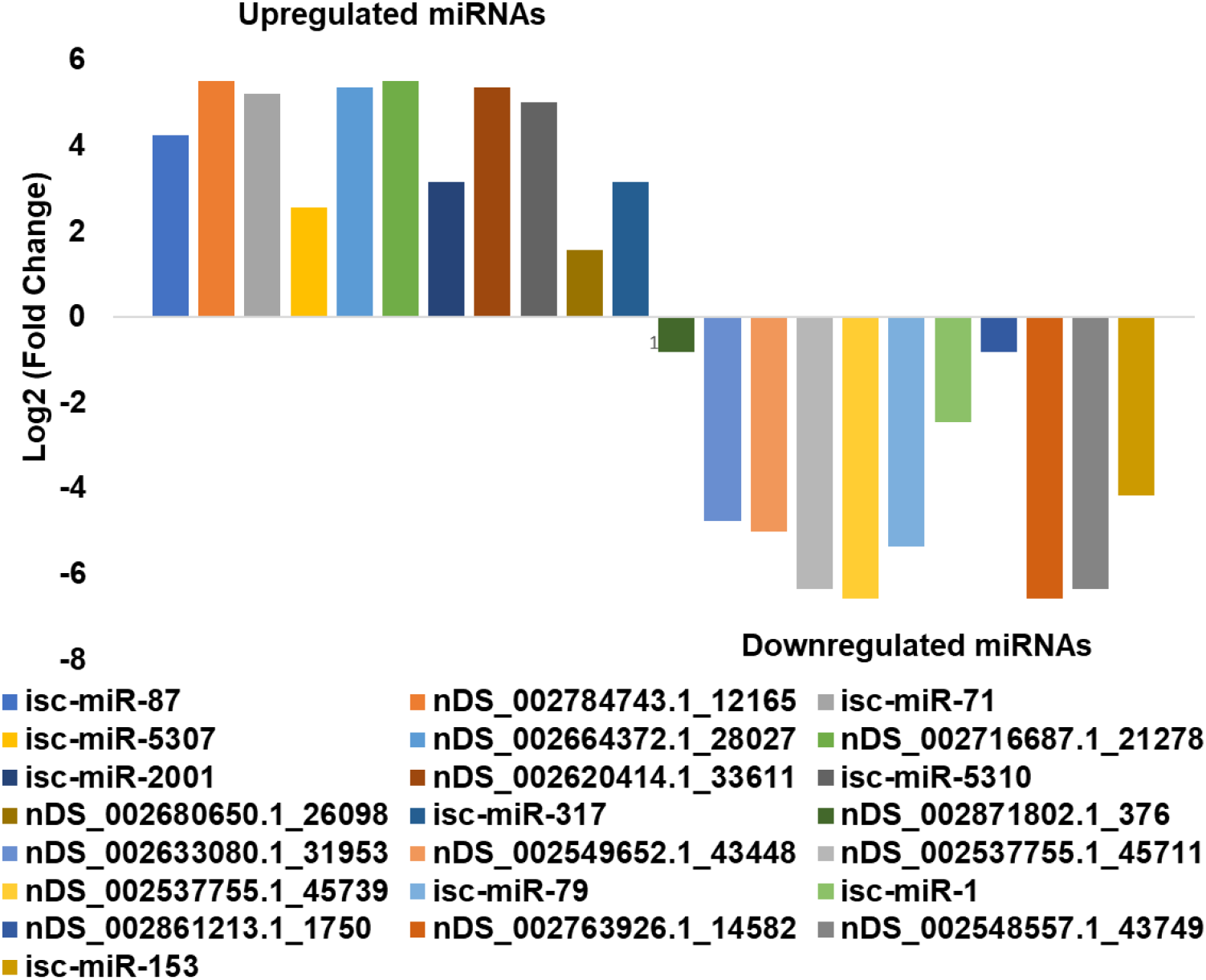
*In silico* differential expression of predicted miRNAs in *Borrelia burgdorferi-*infected, partially fed salivary glands relative to partially fed clean salivary glands. EdgeR was used for differential expression analysis. 12 predicted miRNAs were down-regulated, 11 were upregulated, while 39 were unaffected. miRNAs with a log2 fold-change expression > |1| and FDR ≤ 0.1 were considered significantly differentially expressed (see Table 1). Also, in Table 2, putative roles and targets of these DE conserved miRNAs have been provided based on available studies in other arthropods.

**Table 1.** *In silico* differential expression of *Ixodes scapularis* miRNAs in salivary glands after *Borrelia burgdorferi* infection compared with uninfected salivary glands. miRNAs with a log2 fold-change expression > |1| and FDR ≤ 0.1 were considered significantly differentially expressed. Values highlighted in **red** indicate significant upregulation and values highlighted in **green** indicate significant downregulation.

| Differentially expressed miRNA | Log<sub>2</sub>FC | logCPM | LR | P-value | FDR |
| --- | --- | --- | --- | --- | --- |
| **isc-miR-87** | **4.245604** | 11.83289 | 33.20241 | 8.30E-09 | 4.65E-08 |
| **nDS_002784743.1_12165** | **5.511035** | 11.48887 | 29.7299 | 4.97E-08 | 1.55E-07 |
| **isc-miR-71** | **5.226443** | 11.95849 | 48.94389 | 2.63E-12 | 3.69E-11 |
| **isc-miR-5307** | **2.570363** | 12.34313 | 24.71399 | 6.65E-07 | 1.55E-06 |
| **nDS_002664372.1_28027** | **5.362184** | 11.43227 | 26.80077 | 2.26E-07 | 5.84E-07 |
| **nDS_002716687.1_21278** | **5.511153** | 11.48887 | 29.72059 | 4.99E-08 | 1.55E-07 |
| **isc-miR-2001** | **3.146271** | 11.78845 | 19.75379 | 8.81E-06 | 1.64E-05 |
| **nDS_002620414.1_33611** | **5.361301** | 11.43227 | 26.76848 | 2.29E-07 | 5.84E-07 |
| **isc-miR-5310** | **5.009416** | 11.31195 | 20.86089 | 4.94E-06 | 1.06E-05 |
| **nDS_002680650.1_26098** | **1.567661** | 15.43032 | 103.3589 | 2.80E-24 | 7.83E-23 |
| **isc-miR-317** | **3.146271** | 11.78845 | 19.75379 | 8.81E-06 | 1.64E-05 |
| **nDS_002871802.1_376** | **-0.80359** | 15.73883 | 19.22785 | 1.16E-05 | 1.91E-05 |
| **nDS_002633080.1_31953** | **-4.7431** | 11.95812 | 10.11663 | 0.001469 | 0.001959 |
| **nDS_002549652.1_43448** | **-4.9974** | 12.11011 | 12.13051 | 0.000496 | 0.000731 |
| **nDS_002537755.1_45711** | **-6.34111** | 13.07343 | 30.24285 | 3.81E-08 | 1.52E-07 |
| **nDS_002537755.1_45739** | **-6.57147** | 13.2637 | 36.56262 | 1.48E-09 | 1.03E-08 |
| **isc-miR-79 dme-miR-9a-5p** | **-5.35587** | 12.34273 | 15.64359 | 7.65E-05 | 0.000119 |
| **isc-miR-1 dme-miR-1-3p** | **-2.44072** | 12.67077 | 10.50794 | 0.001189 | 0.001664 |
| **nDS_002861213.1_1750** | **-0.80359** | 15.73883 | 19.22785 | 1.16E-05 | 1.91E-05 |
| **nDS_002763926.1_14582** | **-6.57147** | 13.2637 | 36.56262 | 1.48E-09 | 1.03E-08 |
| **nDS_002548557.1_43749** | **-6.34111** | 13.07343 | 30.24285 | 3.81E-08 | 1.52E-07 |
| **isc-miR-153** | **-4.15091** | 11.64621 | 6.563386 | 0.01041 | 0.013249 |

**Table 2.** List of differentially expressed microRNAs detected in *Borrelia burgdorferi* infected tick salivary glands and their putative roles

| microRNA | Role of ortholog microRNA | Target genes | Reference |
| --- | --- | --- | --- |
| isc-miR-153 | Development, immune response |  | [82] |
| isc-miR-1 | Stress response, immunity, development, facilitating infection | *HSP60, HSP70, GATA4* | [49, 50, 51, 52, 83, 84, 85, 86, 87] |
| isc-miR-79 | Innate immunity, differentiation, apoptosis | *Roundabout protein 2 pathway (robo2), DRAPER, HP2 (Hemolymph protease), P38, pvr, puc* | [55, 56, 57, 88, 89, 90, 91] |
| isc-miR-317 | Negative influencer of Toll pathway, host homeostatic response (targeting Gap junctions, TRP channels), mitotic regulation, Fecundity | *Dif-Rc, Thioredoxin peroxidase-2, STAT* | [45, 91, 92, 93, 94] |
| isc-miR-2001 | Under extreme environmental conditions |  | [95] |
| isc-miR-5307 | Propagation of Powassan virus inside vero cells |  | [96] |
| isc-miR-71 | Stress response, development, facilitates WSSV infection | *cap-1, riok1, Actin1, myD88, IML3* | [91, 97, 98, 99, 100] |
| isc-miR-87 | Pathogen survival (disruption of Toll and IMD pathway) | *Serine/Threonine Kinase, Toll 1A, Putative TLR 5b, FADD* | [60, 62, 63, 91, 101, 102] |
| isc-miR-5310 | Pathogen survival by modulating signaling pathways, Feeding behavior |  | [64-68, 81, 103] |

miR-2001 was upregulated in *B. burgdorferi*-infected salivary glands compared with controls. miR-2001 is not present in the *Drosophila* genome but is an evolutionarily conserved miRNA in ticks [44]. It has previously been shown to play a role in host immunomodulation required for pathogen survival. miR-2001 has been detected in *I. ricinus* tick saliva [45] and *H. longicornis* tick saliva extracellular vesicles (EVs) [46] along with several other miRNAs, suggesting that EVs containing these miRNAs could be transferred to the host to modulate host cellular functions to facilitate tick and pathogen survival [45, 46]. EVs are involved in the intercellular transfer of miRNAs, lipids, and proteins and the disposal of unnecessary cell contents [46, 47]. The discovery of EVs in the excretory-secretory products of ectoparasites suggests that EVs are probably taken up by host cells, deliver their cargoes to the host, and favor immunomodulation, pathogen survival, and disease progression [48, 80]

miR-1 was downregulated in *B. burgdorferi*-infected tick salivary glands compared with uninfected salivary glands. miR-1 belongs to a family of miRNAs including miR-7 and miR-34 conserved across fruit flies, shrimps, and humans, where it modulates similar pathways (development, apoptosis) and is upregulated during stress insults [49]. In mosquitoes, miR-1 is upregulated during *Plasmodium* infection [50] and also facilitates or promotes West Nile virus infection [51]. miR-1 has been shown to be generally upregulated in response to infection, while we detected its downregulation in *B. burgdorferi*-infected SGs. In *Listeria*-infected macrophages, miR-1 promotes IFN-γ-dependent activation of the innate immune response [52].

Tick saliva and salivary gland extracts reduce IFN-γ and IL-2 production in T cells and inhibited T cell proliferation [53, 54] suggesting immune suppression, a possible survival mechanism for tick pathogens. mir-79 was also downregulated in *Borrelia burgdorferi* infected salivary glands, and mir-79 has been shown to participate in immunity and other processes such as cellular differentiation, neurogenesis, and apoptosis. In *Rhipicephalus haemaphysaloides*, mir- 79 was downregulated upon lipopolysaccharide (LPS) induction in female and male ticks, suggesting a role for mir-79 in LPS-mediated stimulation of the innate immune response [55]. The JNK pathway is an immune response pathway against Gram-negative bacteria [56], and mir- 79 is known to disrupt JNK signaling by targeting its component genes *pvr* (CG8222) and *puc* (CG7850) [57]. Our detection of the downregulation of mir-79 in *B. burgdorferi*-infected salivary glands is probably due to stimulation of the JNK pathway as a tick immune response against *B. burgdorferi*. Although *B. burgdorferi* is described as an atypical Gram-negative bacterium due to its double membrane, it lacks classical lipopolysaccharide (LPS). It also has a different cellular organization and membrane composition to other diderms [58]. However, surprisingly in *A. phagocytophilum* (an intracellular Gram-negative bacterial pathogen)-infected ticks, mir-79 was upregulated to facilitate infection by targeting the Roundabout protein 2 pathway (Robo2) [59], suggesting different roles for mir-79 in ticks when infected with different bacterial pathogens. *Borrelia burgdorferi* (Bb) is extracellular and a Gram-negative bacterial pathogen, but *A. phagocytophilum* is a well-known intracellular a Gram-negative bacterial pathogen. Do extracellular and pseudo-Gram-negative status of *B. burgdorferi* make it a different bacterial pathogen than Gram negative or positive? More clarity of tick immune response is required in case of *B. burgdorferi* infection.

In *Drosophila [35]*, miR-317 negatively regulates Toll pathway signaling, and its upregulation in *B. burgdorferi-*infected salivary glands may suggest a similar role to facilitate *B. burgdorferi* survival inside tick salivary glands. *In silico*, miR-317 targets Dif-Rc, an important transcription factor in the Toll pathway in *Drosophila* [35] and STAT in JAK-STAT signaling in *Manduca sexta [60]*. An *in silico* study in *I. ricinus* suggested a combinatorial effect of tick salivary miRNAs on host genes important for maintaining host homeostasis and tick-host interactions, including miR-317 targeting gap junctions and TRP channels, which play significant roles in host homeostatic responses [45]. miR-71 was upregulated in *B. burgdorferi*-infected salivary glands,and its predicted targets include *MyD88*, which is activated when ligands bind to the Toll-like receptor (TLR), interleukin 1 receptor (IL-1R), or IFN-γR1 and trigger MyD88-mediated signaling and pro-inflammatory cytokine responses. Another miR-71 target is IML3, an arthropod immunolectin that recognizes LPS on Gram-negative bacteria as a part of arthropod immune defenses. Immunolectins are also predicted targets of miR-87, -276, -9a, and -71 [60]. We hypothesize that miR-71 disrupts tick immune pathways and protects *B. burgdorferi* in the salivary gland. It has also been shown to prolong the life and regulate stress responses in nematodes, being upregulated in the Dauer larval stage when food or other life-sustaining resources are scarce [61].

miR-87 was upregulated in *B. burgdorferi-*infected salivary glands. Previous studies in other arthropods such as *Manduca sexta* and *Aedes albopictus* have suggested a role in disrupting innate immunity, particularly via IMD and Toll signaling pathways [60, 62, 63]. In *Aedes albopictus*, miR-87’s predicted targets are Toll pathway signaling Ser/Thr kinase, Toll-like receptor Toll1A, class A scavenger receptor with Ser-protease domain, galectin [63], and TLR5b [62], while in *Manduca sexta* its predicted target is FADD, an adaptor protein involved in DISC formation [60]. Our *in silico* data also showed upregulation of miR-5310 in *B. burgdorferi*-infected salivary glands. miR-5310 is a tick-specific miRNA [44], and a recent study demonstrated its downregulation in *Anaplasma phagocytophilum*-infected nymphs compared with unfed uninfected nymphs [64]. Previous studies have also indicated modulation of signaling events via miR-5310 upon *A. phagocytophilum* infection [64-68, 81] In *B. burgdorferi* infection, we speculate that might modulate signaling events and protect *B. burgdorferi* in the tick salivary glands. miR- 5310 might also be involved in tick feeding, as it was found to be downregulated in *Rhipicephalus microplus* tick larvae upon exposure to host odor but not being allowed to feed [44].

### Prediction of target genes and gene ontology (GO) and functional enrichment analyses of the target network

Target proteins were used to build a high-confidence interaction network (interaction scores >0.9).

STRING web analysis (**Figure 4**) showed that the target proteins of 23 DE miRNAs (11 upregulated and 12 downregulated) had similar interactions to those expected for a random set of proteins of similar size sampled from the *I. scapularis* genome (nodes = 687, edges = 79, average node degree = 0.23, average local clustering coefficient = 0.0966, expected number of edges = 77, PPI enrichment *p*-value = 0.411). This does not necessarily mean that these selected proteins are not biologically meaningful, rather that these tick proteins may not be very well studied and their interactions might not yet be known to STRING.

**Figure 4.**
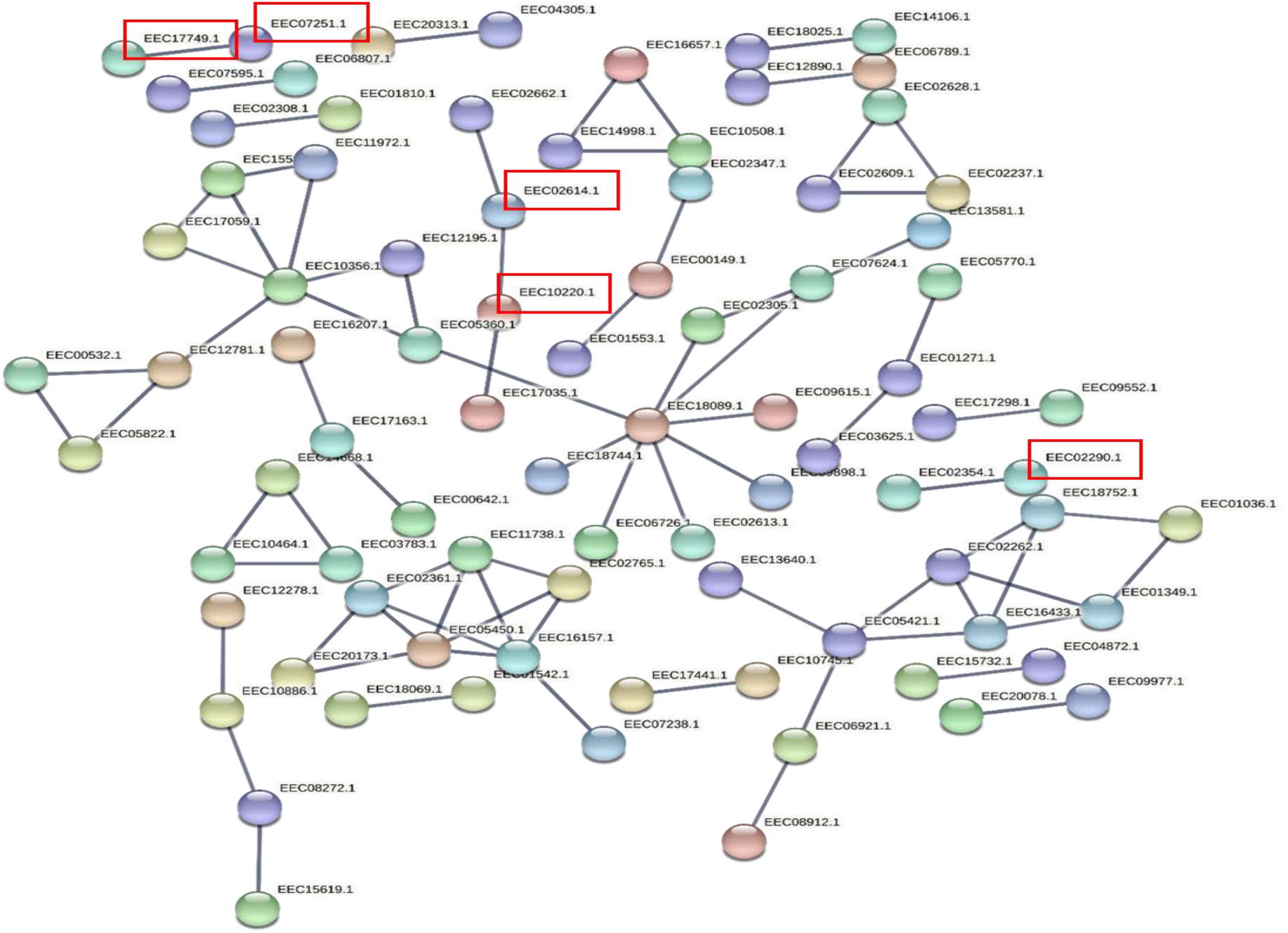
A network built exclusively from *Ixodes scapularis* proteins targeted by *in silico* differentially expressed miRNAs in *Borrelia burgdorferi-*infected, partially-fed salivary glands relative to partially fed clean salivary glands. Red boxes indicates the proteins involved in significant KEGG pathways such as sphingolipid metabolism (EEC07251.1, KEGG:R02541); valine, leucine, and isoleucine degradation (EEC10220.1, KEGG:R04188) ; lipid transport and metabolism (EEC02614.1, KEGG:R01178) ; exosome biogenesis and secretion (EEC07251.1, EEC17749.1, KEGG:R02541) ; and phosphate-containing compound metabolic process (EEC02290.1, KEGG:R00004)/

Many target genes were identified for the DE miRNAs using the sRNAtoolbox miRNAconsTarget program [39]. To minimize false-positive targets, we chose only those targets predicted by all three miRNA target prediction algorithms (TargetSpy, MIRANDA, and PITA). Forty-one KEGG pathways were enriched for target genes (proteins) of DE miRNAs (**Supplementary Table S2**) and included sphingolipid metabolism; valine, leucine, and isoleucine degradation; lipid transport and metabolism; exosome biogenesis and secretion; and phosphate-containing compound metabolic process (**Figure 4**). Gene ontology (GO) analysis indicated that most target genes of DE miRNAs play significant roles in cellular processes, metabolic processes, biological regulation, developmental processes, and responses to stimuli (**Figure 5**). Surprisingly, immune response genes were one of the least affected functions.

**Figure 5.**
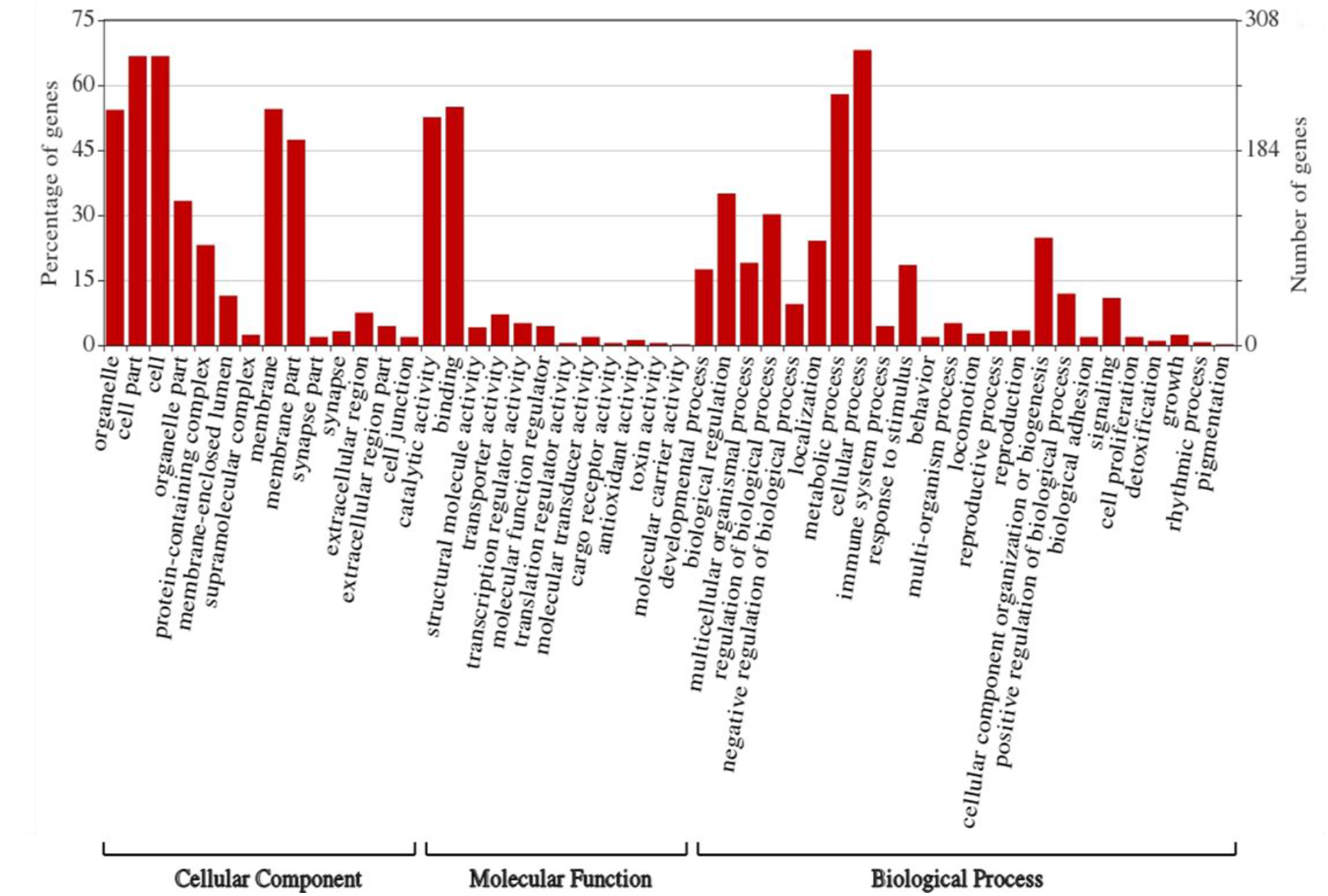
Gene ontology (GO)-derived biological processes related to genes targeted by differentially expressed miRNAs in partially fed *B. burgdorferi*-infected salivary glands relative to partially fed uninfected salivary glands from *Ixodes scapularis* ticks.

Lipid metabolism was one of the main KEGG pathways detected by Pannzer and STRING analyses and was predicted to be regulated by tick salivary gland miRNAs miR-1, miR-5310, miR- 71, and miR-79. It has previously been shown that binding of *B. burgdorferi* to host glycosphingolipid can contribute to tissue-specific adhesion of *B. burgdorferi*, and the inflammatory process in Lyme borreliosis might be affected by interactions between *B. burgdorferi* and glycosphingolipid [69]. Therefore, we hypothesize that tick miRNAs (via saliva) promote sphingolipid synthesis inside hosts to promote *Borrelia* adhesion, and indeed there is evidence that its infection affects lipid metabolism in hosts [70].

### Validation of DE miRNAs by qRT-PCR

The expression of DE miRNAs were validated in *B. burgdorferi*-infected and uninfected ISE6 cells by qRT-PCR (**Figure 6; Supplementary Table S3**), which closely mirrored the RNA-seq data for many targets, although some differences were not statistically significant by qRT-PCR and isc- miR-317 was downregulated rather than upregulated. These differences could be due to the use of different methodologies to quantify miRNA expression [10].

**Figure 6.**
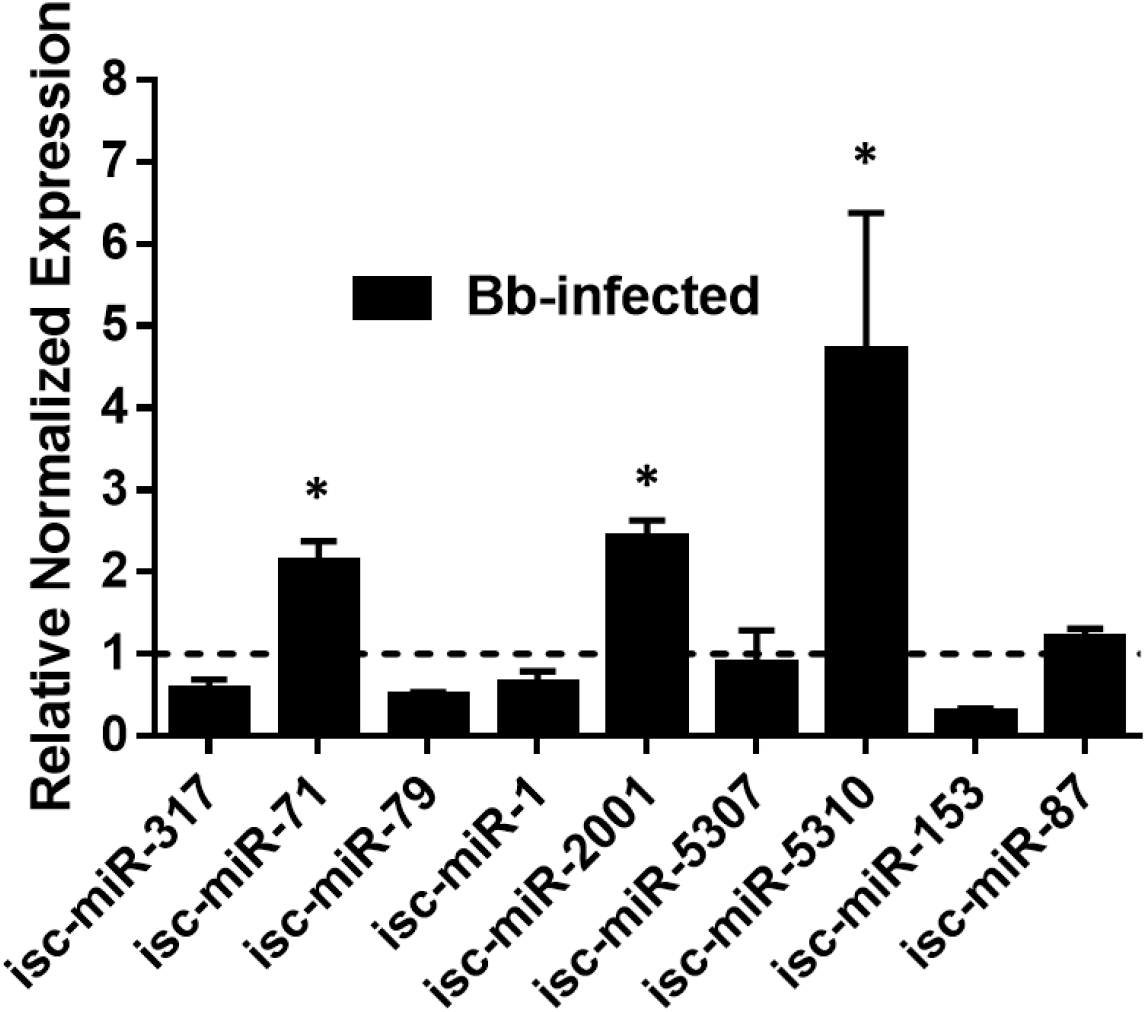
qRT-PCR validation of differentially expressed miRNAs detected in *Borrelia burgdorferi-*infected, partially-fed salivary glands relative to partially-fed uninfected salivary glands from *Ixodes scapularis* ticks. qPCR validation was performed in *Borrelia burgdorferi* infected and clean ISE6 cells. Expression of miRNAs was normalized to clean ISE6 cells (indicated as 1 on the y-axis). Statistical significance for qRT-PCR-based differential expression was determined by the 2-tailed Student’s *t*-test, where * is *p*<0.05.

## Conclusions

This is the first comprehensive miRNA profiling study of *Ixodes scapularis* salivary glands with and without *Borrelia burgdorferi* infection. Here we identified several potential miRNAs targets in tick salivary glands which might play a significant role in *Borrelia* colonization, survival, transmission, and host immunomodulation. Functional validation of these miRNAs is now required. Further characterization of tick salivary gland miRNAs would contribute to a better understanding of the mechanisms underpinning *Borrelia* transmission and propagation inside hosts, not least due to its special status as an extracellular spirochaete and atypical Gram-negative organism that might exploit different survival mechanisms. The impact of *B. burgdorferi* on miRNA expression must also be studied in other tick tissues and hosts to understand cues of its vector competence in ticks and immunomodulation in vertebrates.

