## Supplementary material for "Identification of microRNAs in the Lyme disease vector *Ixodes scapularis*": Fig.1S, Table 1S, Table 2S, Table 3S

**Supplementary Materials**

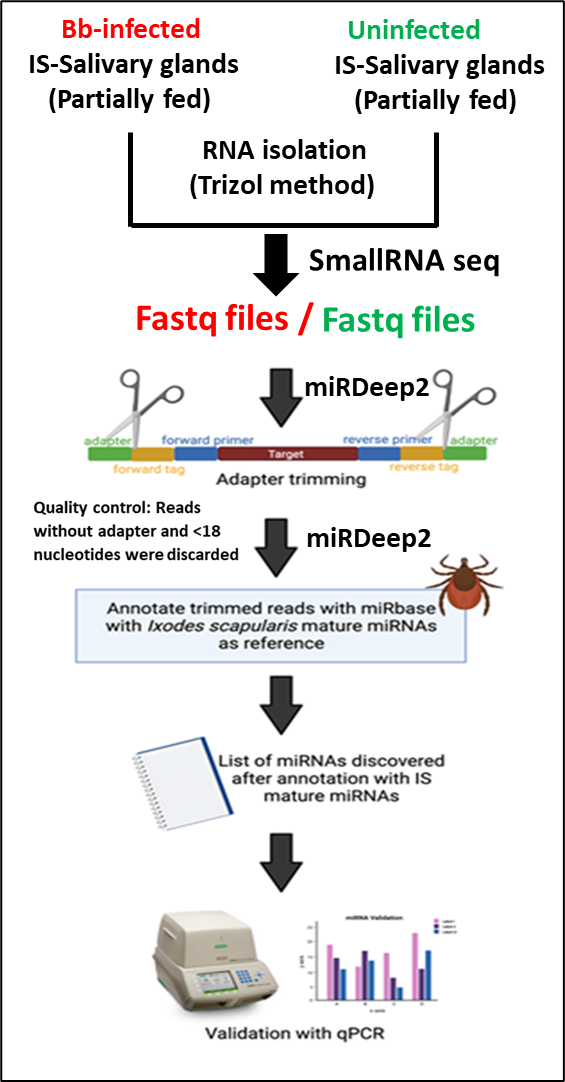

**Figure S1.** Schematic of the experimental plan and data analysis.

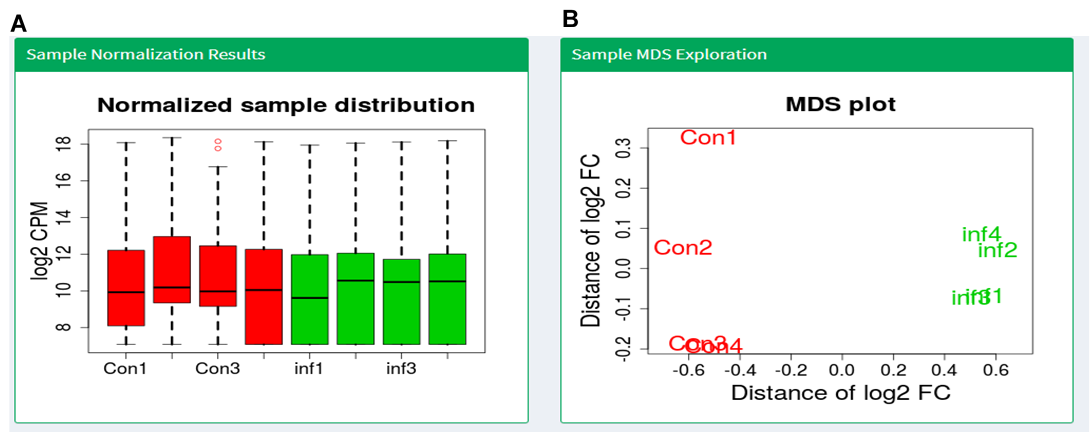

**Figure S2. A.** Normalized sample distribution for *B. burgdorferi*-infected (inf1, inf2, inf3, and inf4) and uninfected (Con1, Con2, Con3, and Con4) salivary glands. **B.** Multidimensional ccaling (MDS) plot shows variation in differential expression among uninfected samples (Con1, Con2, Con3, and Con4), where the distance between sample labels indicates dissimilarity in log2-fold change (FC).

### **Table S1A. MicroRNA profiling of *Ixodes scapularis* salivary glands (novel miRNAs predicted by miRDeep2).**

| **Provisional id** | **miRDeep2 Score** | **mature read count** | **miRbase miRNA with same seed** | **consensus mature sequence** |
| --- | --- | --- | --- | --- |
| isc-miR-4968-5p | 1.80E+04 | 36306 | dme-miR-4968-5p | aagcugcccaaugaagggcug |
| NW_002532706.1_43464  isc- miR-92a-3p | 1.50E+04 | 29572 | dme-miR-92a-3p | uauugcacucgucccggccuuu |
| NW_002871802.1_367  isc -miR-125-5p | 1.20E+04 | 12052 | dme-miR-125-5p | ucccugagacccuaacuuguga |
| NW_002692816.1_23152  isc-miR-6-3p | 7.60E+03 | 10940 | dme-miR-6-3p | uaucacagccaucuuugaugacc |
| nDs_002867181.1_816 | 2.40E+03 | 4514 |  | gccaacccacuaacugacggga |
| nDs_002653147.1_27921 | 2.30E+03 | 4514 |  | gccaacccacuaacugacggga |
| nDs_002794895.1_10282 | 2.30E+03 | 4514 |  | gccaacccacuaacugacggga |
| nDs_002645436.1_28906 | 1.80E+03 | 3392 |  | gcuguuaguuugugggguggug |
| nDs_002860940.1_1699 | 1.80E+03 | 3392 |  | gcuguuaguuugugggguggug |
| nDs_002508325.1_46074 | 1.80E+03 | 3279 |  | gcuguuaguuugugggguggug |
| nDs_002699119.1_22404 | 1.70E+03 | 3392 |  | gcuguuaguuugugggguggug |
| NW_002767753.1_13364  isc-miR-958-3p | 1.70E+03 | 3446 | dme-miR-958-3p | ugagauucaacuccuccaacuucu |
| nDs_002667887.1_26137 | 1.70E+03 | 3392 |  | gcuguuaguuugugggguggug |
| nDs_002810888.1_8213 | 1.70E+03 | 3392 |  | gcuguuaguuugugggguggug |
| nDs_002642603.1_29251 | 1.70E+03 | 3392 |  | gcuguuaguuugugggguggug |
| nDs_002815928.1_7504 | 1.00E+03 | 2109 |  | caagaagauagaaucaaugaga |
| nDs_002861213.1_1662 | 9.50E+02 | 1855 |  | cuugggcuggaguucgucguugu |
| isc-miR-190-5p | 6.60E+02 | 621 | dme-miR-190-5p | agauauguuugauauucuugguu |
| nDs_002614457.1_32408 | 6.30E+02 | 1205 |  | cacgucgaggcuucagguugu |
| nDs_002614457.1_32404 | 5.20E+02 | 844 |  | uacgucgagaccucagguuga |
| nDs_002867165.1_832 | 5.00E+02 | 994 |  | ucguuuucgagaucgugaaccu |
| nDs_002728031.1_18707 | 4.90E+02 | 952 |  | aguggucaugucuucgcacugga |
| nDs_002621331.1_31664 | 4.50E+02 | 812 |  | gugugagaaaaugguuggcaca |
| nDs_002614457.1_32406 | 4.50E+02 | 857 |  | cacgucgagauuucaagcuaug |
| nDs_002735932.1_17756 | 4.10E+02 | 752 |  | ugaggucaugcucggugccuug |
| nDs_002873519.1_126 | 3.70E+02 | 733 |  | auaucuuuuuuuaacaucggcu |
| nDs_002784743.1_11607 | 2.10E+02 | 361 |  | cauucuugcagcgaaggguugu |
| nDs_002620414.1_31747 | 1.50E+02 | 213 |  | cagugcuucugcagugcaggc |
| nDs_002871485.1_401 | 1.40E+02 | 280 |  | acagaaucgagcgcuagggcga |
| nDs_002668420.1_26105 | 1.10E+02 | 200 |  | ggcacaaggcugucggaugcga |
| nDs_002680650.1_24606 | 1.00E+02 | 146 |  | cucagaaaugcuaggcuaucgg |
| NW_002527183.1_43988  isc-miR-263b-5p | 1.00E+02 | 184 | dme-miR-263b-5p | cuuggcacugaaagaauucaca |
| nDs_002685244.1_24008 | 9.90E+01 | 185 |  | cuuguuugggcaaaugggugac |
| nDs_002814339.1_7692 | 9.60E+01 | 185 |  | cacgggccccagcugaacgcu |
| nDs_002737590.1_17488 | 9.00E+01 | 170 |  | cagucaguggcuuugguucauc |
| nDs_002737673.1_17464 | 8.60E+01 | 173 |  | guacacguucgggcuuccaccc |
| nDs_002564688.1_39220 | 7.40E+01 | 96 |  | gguuauuuucagcggcugucgu |
| nDs_002871133.1_436 | 7.00E+01 | 146 |  | agccauuuuuugaagucgaca |
| nDs_002737711.1_17449 | 4.80E+01 | 84 |  | agagugacgcagagaacaaauc |
| nDs_002675644.1_25181 | 4.50E+01 | 84 |  | aaaaggacagaacauguagaca |
| nDs_002763926.1_13886 | 3.90E+01 | 69 |  | uguagccggauugugggacugg |
| nDs_002531612.1_43580 | 3.70E+01 | 53 |  | cacugugcgcgauugguucaac |
| nDs_002722741.1_19330 | 3.30E+01 | 75 |  | auugaacgagaggacauaguc |
| nDs_002567548.1_38945 | 3.10E+01 | 52 |  | ucaguguccgucgccggaagg |
| NW_002787101.1_11264  isc-miR-31b-5p | 3.10E+01 | 26 | dme-miR-31b-5p | aggcaagaagucuuuuaggaug |
| nDs_002754008.1_15271 | 3.00E+01 | 70 |  | ucacuuuacgucguaguacaag |
| nDs_002538194.1_42559 | 3.00E+01 | 53 |  | uuaguacgauccgucgaaagac |
| nDs_002801682.1_9417 | 3.00E+01 | 54 |  | uguagcaugcuccaagcggagg |
| nDs_002508917.1_46002 | 3.00E+01 | 54 |  | uguagcaugcuccaagcggagg |
| nDs_002568676.1_38752 | 2.70E+01 | 48 |  | aaggagacaaggaaacaaacacc |
| nDs_002642725.1_29244 | 2.60E+01 | 58 |  | acaacccggaaucgucauggcc |
| nDs_002670695.1_25779 | 2.60E+01 | 40 |  | gcuggaucguaaaacuggugcca |
| nDs_002721909.1_19420 | 2.40E+01 | 44 |  | gagaggucgguagaguugcgc |
| nDs_002548557.1_41255 | 2.40E+01 | 40 |  | cuagugugacugucacagugg |
| nDs_002856865.1_2668 | 2.40E+01 | 40 |  | cccggcgagaaaucuggcggccg |
| nDs_002588255.1_36295 | 2.10E+01 | 24 |  | agacacgcuacugacuaucgucg |
| nDs_002591771.1_35729 | 1.80E+01 | 12 |  | cccgguacggacugcgcauguc |
| NW_002637235.1_29734  isc-miR-4987-5p | 1.80E+01 | 20 | dme-miR-4987-5p | augcaacagaggcgagaugagacg |
| nDs_002730915.1_18372 | 1.80E+01 | 21 |  | gaggaagaaaacgcgaaaggac |
| nDs_002527216.1_43980 | 1.70E+01 | 26 |  | aggugaaugaauacccagucgaug |
| nDs_002829027.1_5932 | 1.60E+01 | 15 |  | acgaaaucccgcggaucgcauug |
| nDs_002741178.1_17075 | 1.60E+01 | 27 |  | uugggcuuuaaacaucaugagc |
| nDs_002718594.1_19992 | 1.50E+01 | 25 |  | ucacaggacgugcaacggugcu |
| nDs_002718594.1_19994 | 1.50E+01 | 25 |  | ucacaggacgugcaacggugcu |
| NW_002681830.1_24464  isc-bantam-3p | 1.40E+01 | 17 | dme-bantam-3p | cgagaucauacgucgguggugg |
| nDs_002515969.1_45216 | 1.40E+01 | 17 |  | acaauugaaguggaucccgaau |
| nDs_002770056.1_13170 | 1.40E+01 | 24 |  | aacgaaucgaaggucaaagugc |
| nDs_002635366.1_29915 | 1.30E+01 | 36 |  | cuugugaugcucucguucgacc |
| nDs_002745696.1_16495 | 1.20E+01 | 16 |  | uccgaggaaguguccugacaga |
| nDs_002633080.1_30207 | 1.10E+01 | 17 |  | uauuccuuccuuuuguccgguc |
| nDs_002862093.1_1546 | 1.00E+01 | 18 |  | cuauuggacgaaaaggauagcu |
| nDs_002770523.1_13144 | 8.5 | 11 |  | ugaagaucuccaacuuggccucg |
| nDs_002637837.1_29679 | 8 | 10 |  | aaagaaauacggacgacaggag |
| NW_002716687.1_20192  isc-miR-5-5p | 5.7 | 209 | dme-miR-5-5p | caaggaaccgacgaaucgagugu |
| NW_002522525.1_44526  isc-miR-1007-5p | 5.6 | 30 | dme-miR-1007-5p | ucaguguuuggcuggaacccg |
| NW_002784743.1_11604  isc-miR-277-3p | 5.4 | 1487 | dme-miR-277-3p | uaaaugcauuaucuggaaug |
| NW_002788893.1_11074  isc-miR-4957-3p | 5.3 | 52 | dme-miR-4957-3p | ucagcugcgugaacgcgggcgcc |
| NW_002799140.1_9764  isc-miR-2a-1-5p | 4.5 | 101 | dme-miR-2a-1-5p | uucucaaauguccgccacucg |
| nDs_002745459.1_16501 | 2.7 | 25 |  | accccggggguggucacgugac |
| nDs_002813294.1_7883 | 2.5 | 19 |  | acggagcgacugaacaagaca |
| nDs_002829114.1_5911 | 2.4 | 19 |  | agagaaaaagacagguagaacg |
| nDs_002759900.1_14325 | 2.4 | 95 |  | uugccgccgucguucauggaugg |
| nDs_002815075.1_7597 | 2.4 | 98 |  | uugccgccgucguucauggaugg |
| nDs_002513235.1_45498 | 2.3 | 26 |  | ccucgguuuucggcuggcacucg |
| nDs_002676299.1_25117 | 2.3 | 22 |  | ccaccgguuacgcucugcgcc |
| nDs_002700913.1_22252 | 2.3 | 123 |  | gggacgaaacagacgacacagc |
| nDs_002684507.1_24073 | 2.3 | 258 |  | caggcgggagaacaauguccu |
| nDs_002756339.1_14956 | 2.3 | 13 |  | caagucgcggcugcacgggcgcc |
| nDs_002774760.1_12785 | 2.3 | 12 |  | cuucucucugcugccguggccc |
| nDs_002696292.1_22736 | 2.3 | 39 |  | cugccucgcgcuggucuccgcu |
| nDs_002550131.1_40913 | 2.2 | 19 |  | cgccucagcuguuucugcucc |
| nDs_002542368.1_42013 | 2.2 | 18 |  | cacgucacagcguccgcucaag |
| nDs_002586787.1_36439 | 2.2 | 37 |  | uuggacgaaaaaaagagccgacu |
| nDs_002607638.1_33547 | 2.2 | 17 |  | cuagccaaucguggcggucggc |
| nDs_002836532.1_4888 | 2.2 | 22 |  | ccaccgguuacgcucugcgcc |
| nDs_002597262.1_35126 | 2.1 | 11 |  | cgggaucucggaggccaugaa |
| nDs_002662116.1_26803 | 2.1 | 21 |  | caggauccugacgcuggugggcaca |
| nDs_002636820.1_29767 | 2.1 | 7545 |  | caaggcaaugaacaugaucuca |
| nDs_002799981.1_9646 | 2.1 | 24 |  | uccguccguuccgucccucu |
| nDs_002612646.1_32606 | 2.1 | 7545 |  | caaggcaaugaacaugaucuca |
| nDs_002629651.1_30593 | 2.1 | 819 |  | cuucguagucggauuaugacu |
| nDs_002601819.1_34393 | 2.1 | 32 |  | ucgaccgcucgccacggcugg |
| nDs_002540798.1_42197 | 2.1 | 20 |  | caugucacugcguccgcgcaag |
| nDs_002654127.1_27778 | 2 | 200 |  | uuuccgcugucgcuuguagaug |
| nDs_002860687.1_1738 | 2 | 15 |  | gcagaucucagguguagugauc |
| nDs_002665175.1_26443 | 2 | 217 |  | aacuucgugcugcaggagcgccu |
| nDs_002751908.1_15689 | 2 | 49 |  | uaauuggucuuugugagugcuu |
| nDs_002543222.1_41886 | 2 | 149 |  | gccggugucgaucuugaaguuc |
| nDs_002569324.1_38643 | 2 | 22 |  | ccaccgguuacgcucugcgcc |
| nDs_002590536.1_35849 | 2 | 22 |  | uucgguugugccgcuugccu |
| nDs_002601819.1_34394 | 2 | 32 |  | ucgaccgcucgccacggcugg |
| nDs_002792777.1_10524 | 2 | 15 |  | ugagaucucggaggucgugcgcc |
| nDs_002714141.1_20473 | 2 | 83 |  | ggcuucguagucggauuauga |
| nDs_002744939.1_16574 | 1.9 | 829 |  | cuucgaaaucggauuaugacu |
| nDs_002533105.1_43443 | 1.9 | 7545 |  | caaggcaaugaacaugaucuca |
| nDs_002753934.1_15280 | 1.9 | 1842 |  | cauccgguccuaagaagucgaa |
| nDs_002692659.1_23173 | 1.9 | 11 |  | cugccgaaguagcgucugcucu |
| nDs_002733141.1_18085 | 1.9 | 7545 |  | caaggcaaugaacaugaucuca |
| nDs_002816672.1_7394 | 1.9 | 200 |  | uuuccgcugucgcuuguagaug |
| nDs_002738551.1_17311 | 1.9 | 77 |  | cgcgcggacgcugugacgcagc |
| nDs_002523780.1_44319 | 1.9 | 7545 |  | caaggcaaugaacaugaucuca |
| nDs_002828015.1_6065 | 1.9 | 2093 |  | ccaggauuugaacucugggccu |
| nDs_002829464.1_5811 | 1.9 | 7545 |  | caaggcaaugaacaugaucuca |
| nDs_002545827.1_41513 | 1.9 | 44 |  | aagggaaccgugcgagagcugcu |
| nDs_002692659.1_23171 | 1.8 | 11 |  | cugccgaaguagcgucugcucu |
| nDs_002661855.1_26902 | 1.8 | 70 |  | gacauaguaaggauugacg |
| nDs_002765967.1_13675 | 1.8 | 17 |  | uauuccuuccuuuuguccgguc |
| nDs_002574390.1_37989 | 1.8 | 259 |  | uuccgccgucgaucguagaug |
| nDs_002550131.1_40914 | 1.8 | 19 |  | cgccucagcuguuucugcucc |
| nDs_002871802.1_364 | 1.8 | 103 |  | agggccugagaaucuaaccugg |
| nDs_002673151.1_25529 | 1.8 | 29 |  | gcgcacagagagacgaagacug |
| nDs_002809300.1_8415 | 1.8 | 1873 |  | cauccgguccuaagaagucgaa |
| nDs_002738154.1_17397 | 1.8 | 35 |  | cggaaggauaguggguggaccug |
| nDs_002532161.1_43535 | 1.8 | 1943 |  | cauccgguccuaagaagucgaa |
| nDs_002798374.1_9856 | 1.8 | 271 |  | uucgagcgcuaggacagaggccg |
| nDs_002724962.1_19022 | 1.8 | 339 |  | agaaagugcgucuggcggcg |
| nDs_002692659.1_23176 | 1.8 | 108 |  | acuaaaaaacaacggacaagu |
| nDs_002744911.1_16578 | 1.8 | 11 |  | auucggacauccgcaggacguc |
| nDs_002545470.1_41626 | 1.8 | 11 |  | auucggacauccgcaggacguc |
| nDs_002785931.1_11413 | 1.8 | 7545 |  | caaggcaaugaacaugaucuca |
| nDs_002685244.1_24007 | 1.8 | 185 |  | cuuguuugggcaaaugggugac |
| nDs_002784743.1_11583 | 1.7 | 12 |  | aggacgauugagcucgac |
| nDs_002587969.1_36333 | 1.7 | 53 |  | cacgauugccaaacuguaugcu |
| nDs_002661304.1_26957 | 1.7 | 39 |  | aguccgucgucgaucuagccu |
| nDs_002707270.1_21452 | 1.7 | 26 |  | uucgcguucggacguguucgaa |
| nDs_002549652.1_41058 | 1.7 | 1892 |  | caugccguggcuugacca |
| nDs_002737099.1_17587 | 1.7 | 379 |  | uuuggcaggcuuagaaucacuc |
| nDs_002511156.1_45736 | 1.7 | 95 |  | uuugcaccgucgcucguagaug |
| nDs_002608667.1_33381 | 1.7 | 30 |  | auggacuugauguuuuaggcucagu |
| nDs_002699570.1_22364 | 1.7 | 108 |  | acuaaaaaacaacggacaagu |
| nDs_002537755.1_42809 | 1.6 | 53 |  | acaccgguauuuaagagguuaucug |
| nDs_002553320.1_40621 | 1.6 | 11 |  | auucggacauccgcaggacguc |
| nDs_002581904.1_37014 | 1.6 | 12 |  | gccauagcguaacagagugacg |
| nDs_002541749.1_42092 | 1.6 | 1943 |  | cauccgguccuaagaagucgaa |
| nDs_002509761.1_45913 | 1.6 | 74 |  | gacuucagguccaucuggacg |
| nDs_002542368.1_42015 | 1.6 | 18 |  | cacgucacagcguccgcucaag |
| nDs_002650258.1_28240 | 1.6 | 56 |  | gcuguuaguuugugggauggu |
| nDs_002745745.1_16473 | 1.6 | 259 |  | uuccgccgucgaucguagaug |
| nDs_002692816.1_23143 | 1.6 | 105 |  | uuucacagucccuuugacgg |
| nDs_002664372.1_26487 | 1.6 | 18 |  | acuaccgaccgugugugaccg |
| nDs_002677543.1_24907 | 1.6 | 1862 |  | cauccgguccuaagaagucgaa |
| nDs_002505804.1_46391 | 1.6 | 74 |  | gacuucagguccaucuggacg |
| nDs_002855841.1_2713 | 1.6 | 137 |  | acugugauugaacugaacgaca |
| nDs_002592079.1_35704 | 1.6 | 14 |  | gaguagaaugccugaauguugcu |
| nDs_002653831.1_27809 | 1.6 | 114 |  | aaggcgaacgcugaccugggcc |
| nDs_002630829.1_30421 | 1.5 | 17 |  | cgacgaauuucuggagaucgca |
| nDs_002535729.1_43080 | 1.5 | 331 |  | cuucgcucugcuaaaguggacc |
| nDs_002590762.1_35829 | 1.5 | 95 |  | uugccgccgucguucauggaugg |
| nDs_002617786.1_32035 | 1.5 | 18 |  | ucugucccugguuuuggcgcu |
| nDs_002834843.1_5113 | 1.5 | 21 |  | cgagggcuguccagaguucuug |
| nDs_002844434.1_4087 | 1.5 | 258 |  | uuccgccgucgaucguagaug |
| nDs_002838169.1_4671 | 1.5 | 47 |  | accauuugggucucuguccucu |
| nDs_002567550.1_38944 | 1.5 | 97 |  | uuugcaccgucgcucguagaug |
| nDs_002639690.1_29492 | 1.5 | 2928 |  | guuguacccagucgucgauguc |
| nDs_002530398.1_43703 | 1.4 | 31 |  | cagggccuaaaugacauuuacacu |
| nDs_002829616.1_5794 | 1.4 | 15 |  | acggcuggcgucucgcugcacc |
| NW_002626049.1_30948  isc-miR-981-3p | 1.4 | 4936 | dme-miR-981-3p | uucguugucguagaaaccugau |
| nDs_002790353.1_10903 | 1.4 | 20 |  | cacgucacagcguccgcucaag |
| nDs_002721909.1_19415 | 1.4 | 312 |  | caugacugucaucuuugcaucu |
| nDs_002592028.1_35707 | 1.4 | 26 |  | cucgaggucuacccgcucuccu |
| nDs_002697784.1_22512 | 1.4 | 121 |  | uuggacuuugagcauggcgaagg |
| nDs_002515625.1_45241 | 1.3 | 24 |  | augcccacugucccaguuucugc |
| nDs_002815935.1_7502 | 1.3 | 15 |  | ucugcgagacgcucaacgacag |
| nDs_002650258.1_28243 | 1.2 | 23 |  | cucaagaucuucggccagucgc |
| nDs_002663292.1_26641 | 1.2 | 137 |  | acugugauugaacugaacgaca |
| nDs_002744911.1_16577 | 1.2 | 11 |  | auucggacauccgcaggacguc |
| nDs_002827450.1_6124 | 1.2 | 20 |  | cacgucacagcguccgcucaag |
| nDs_002545470.1_41625 | 1.2 | 11 |  | auucggacauccgcaggacguc |
| nDs_002553320.1_40622 | 1.2 | 11 |  | auucggacauccgcaggacguc |
| nDs_002598280.1_35014 | 1.1 | 28 |  | ucaacuccuucagcuggugcug |
| nDs_002756375.1_14945 | 1.1 | 52 |  | uucacgaaucugaacgggaca |
| nDs_002613137.1_32572 | 1.1 | 19 |  | ugugaaggccuccugugcucu |
| nDs_002563762.1_39378 | 1 | 18 |  | aagaauaaacaaagguacggau |
| nDs_002768943.1_13309 | 0.9 | 121 |  | ugacgucaucguagugcugcgu |
| nDs_002546088.1_41480 | 0.8 | 18 |  | aucuuuugacacguuggaucu |
| nDs_002856883.1_2662 | 0.8 | 38 |  | ugugagaugauuguugcauaagcg |
| nDs_002615653.1_32278 | 0.8 | 257 |  | uuccgccgucgaucguagaug |
| nDs_002655152.1_27700 | 0.8 | 321 |  | ugaaagcgaguaccaugacugu |
| nDs_002808312.1_8698 | 0.6 | 59 |  | uuguggacgccuggacaagccu |
| nDs_002784518.1_11647 | 0.5 | 26 |  | aucuuuugacacguuggaucu |
| nDs_002860612.1_1764 | 0.4 | 11 |  | aacaugagcgacgucgcuaccg |
| NW_002847019.1_3755  isc-miR-1000-3p | 0.3 | 49 | dme-miR-1000-3p | augcuggggacacugaaaucc |
| nDs_002585048.1_36632 | 0.3 | 9 |  | cuucguagucggauuauga |
| nDs_002569020.1_38698 | 0.2 | 353 |  | gccggaacagucaucguugcug |
| nDs_002820844.1_6987 | 0.2 | 39 |  | auucggcggcacuuugaacgaa |
| NW_002573167.1_38165  isc-miR-2494-3p | 0 | 79 | dme-miR-2494-3p | uucccaguagucccaggaccug |

**Table S1B. mature miRNAs of *Ixodes scapularis* (already available in miRBase) detected in miRDeep2 analysis**

| **tag id** | **miRDeep2 score** | **mature read count** | **mature miRBase miRNA** | **consensus mature sequence** |
| --- | --- | --- | --- | --- |
| NW_002860020.1_1824 | 6.60E+05 | 1296028 | isc-miR-375 | uuuguucguucggcucgaguua |
| NW_002784743.1_11596 | 3.90E+05 | 773543 | isc-miR-10 | uacccuguagauccgaauuugu |
| NW_002651209.1_28145 | 3.60E+05 | 706969 | isc-miR-2001 | uugugaccguuacaaugggcaug |
| NW_002835324.1_5031 | 1.80E+05 | 356093 | isc-miR-12 | ugaguauuacaucagguacuggu |
| NW_002509761.1_45916 | 1.60E+05 | 327576 | isc-bantam | ugagaucauugugaaagcugauu |
| NW_002505804.1_46394 | 1.60E+05 | 327576 | isc-bantam | ugagaucauugugaaagcugauu |
| NW_002871802.1_369 | 4.90E+04 | 96352 | isc-miR-100 | aacccguagauccgaacuugug |
| NW_002656291.1_27550 | 4.10E+04 | 81041 | isc-miR-3931 | uacuuugagucgguacgaauccu |
| NW_002648701.1_28526 | 3.60E+04 | 70402 | isc-miR-8 | uaauacugucagguaaagauguc |
| NW_002835324.1_5033 | 3.40E+04 | 66800 | isc-miR-5307 | uaaucucauuugguaucucuggg |
| NW_002778756.1_12266 | 2.80E+04 | 54755 | isc-miR-79 | ucuuugguuaucuagcuguauga |
| NW_002527183.1_43986 | 2.20E+04 | 43978 | isc-miR-263a | aauggcacuggaagaauucacgg |
| NW_002527216.1_43977 | 2.20E+04 | 42981 | isc-miR-279 | ugacuagauccacacucaucca |
| NW_002741283.1_17066 | 1.50E+04 | 29423 | isc-miR-276 | uaggaacuucauaccaugcucg |
| NW_002822737.1_6809 | 1.40E+04 | 25686 | isc-miR-87 | gugcccggaacuuugucucagccu |
| NW_002697059.1_22632 | 1.30E+04 | 25752 | isc-miR-184 | uggacggagaacugauaagggc |
| NW_002784743.1_11606 | 9.80E+03 | 19228 | isc-miR-317 | ugaacacagcuggugguauaucagu |
| NW_002838206.1_4665 | 8.70E+03 | 16890 | isc-miR-305 | auuguacuucaucaggugcucugga |
| NW_002692816.1_23154 | 6.90E+03 | 13241 | isc-miR-2b | uaucacagccaccuuugaugagcu |
| NW_002692816.1_23150 | 5.60E+03 | 10571 | isc-miR-2a | uaucacagccagcuuugaugagc |
| NW_002692816.1_23156 | 5.50E+03 | 7193 | isc-miR-71 | ucucacuaccuugucuuuguug |
| NW_002804504.1_9143 | 4.60E+03 | 5941 | isc-miR-307 | ccucacucaguuuggcuguggug |
| NW_002535453.1_43180 | 3.00E+03 | 5915 | isc-miR-153 | uugcauagucacaaaagugaug |
| NW_002764570.1_13794 | 2.80E+03 | 5474 | isc-miR-252b | uuaaguaguagugccgcagguaa |
| NW_002604667.1_33926 | 2.80E+03 | 5463 | isc-miR-315 | uuuugauuguugcucagaaggcg |
| NW_002704625.1_21791 | 1.80E+03 | 3647 | isc-miR-1 | uggaauguaaagaaguauggag |
| NW_002734393.1_17986 | 1.00E+03 | 1901 | isc-miR-278 | ccggaugaauuucucgccuggcc |
| NW_002848014.1_3640 | 6.90E+02 | 1368 | isc-miR-5314 | uauagaugaugucuucaugaug |
| NW_002627241.1_30834 | 6.60E+02 | 1302 | isc-miR-5315 | aacacaacauccggacaagcac |
| NW_002527183.1_43990 | 3.70E+02 | 575 | isc-miR-96 | uuuggcacuagcacauuuuugu |
| NW_002522955.1_44449 | 3.10E+02 | 434 | isc-miR-7 | uggaagacuagugauuuuguuguu |
| NW_002784743.1_11634 | 2.10E+02 | 330 | isc-miR-993 | gaagcucguuucuacagguaucu |
| NW_002805128.1_9069 | 1.00E+02 | 90 | isc-miR-5308 | ucugugcuguggagguaauauau |
| NW_002589297.1_36170 | 9.40E+01 | 76 | isc-miR-5305 | uaaguuaaucuccaagcccaau |
| NW_002733905.1_18011 | 5.3 | 272 | isc-miR-124 | uaaggcacgcggugaaugcc |
| NW_002506863.1_46298 | 5.3 | 1270 | isc-miR-133 | uugguccccuucaaccagcugu |
| NW_002542293.1_42026 | 2.1 | 147 | isc-miR-5310 | uguagucuggcagaaacguc |
| NW_002610401.1_33095 | 2 | 69 | isc-miR-5306 | agaguaucacgugacgcuccu |
| NW_002755260.1_15039 | 2 | 28 | isc-miR-1993 | cauuaugcuaguguucgcgggg |
| NW_002796881.1_10053 | 1.9 | 28 | isc-miR-1993 | cauuaugcuaguguucgcgggg |
| NW_002638804.1_29568 | 1.8 | 693 | isc-miR-5309 | caaugcccauggaaaccccgaa |
| NW_002793850.1_10420 | 1.7 | 743 | isc-miR-5312 | uggcugaacguuguuaugcgu |
| NW_002838206.1_4667 | 1.2 | 17097 | isc-miR-275 | ucagguaccugaaguagcgcgc |
| NW_002784743.1_11605 | 1.1 | 19228 | isc-miR-317 | ugaacacagcuggugguauaucagu |
| NW_002822737.1_6807 | -0.8 | 3011 | isc-miR-87 | gugagcaaaguuucaggugugu |
| NW_002796881.1_10054 | -2.2 | 28 | isc-miR-1993 | cauuaugcuaguguucgcgggg |
| NW_002741199.1_17073 | -5.1 | 12 | isc-miR-137 | uuauugcuugagaauacacgu |

**Table S2. KEGG pathways targeted by differentially expressed miRNAs.**

| **Target proteins** | **KEGG ID** | **KEGG pathways** |
| --- | --- | --- |
| EEC07251.1 | KEGG:R02541 | RAS-related protein, putative, Sphingolipid metabolism pathway |
| EEC06395.1 | KEGG:R04030 | Ubiquinone and other terpenoid-quinone biosynthesis, |
|  |  | Metabolic pathways |
|  |  | Biosynthesis of secondary metabolites |
|  |  | Biosynthesis of cofactors |
| EEC17038.1 | KEGG:R02541 |  |
| EEC08267.1 | KEGG:R00253 | Arginine biosynthesis |
|  |  | Alanine, aspartate and glutamate metabolism |
|  |  | Glyoxylate and dicarboxylate metabolism |
|  |  | Nitrogen metabolism |
|  |  | Metabolic pathways |
|  |  | Microbial metabolism in diverse environments |
|  |  | Biosynthesis of amino acids |
| EEC06395.1 | KEGG:R04030 | Fatty-acyl-CoA Synthase (Fatty acid biosynthesis pathway) |
| EEC17038.1 | KEGG:R02541 | RAB-33, putative [Ixodes scapularis], exosome biogenesis and secretion between arthropods and mammals. |
| EEC08267.1 | KEGG:R00253 |  |
| EEC10220.1 | KEGG:R04188 | 4-aminobutyrate aminotransferase, Valine, leucine and isoleucine degradation |
| EEC02614.1 | KEGG:R01178 | medium-chain acyl-CoA dehydrogenase, putative, Lipid transport and metabolism |
| EEC07988.1 | KEGG:R05982 KEGG:R06722 | alpha-mannosidase putative, Protein processing in endoplasmic reticulum, Various types of N-glycan biosynthesis |
| EEC02307.1 | KEGG:R01178 | medium-chain acyl-CoA dehydrogenase, putative, |
| EEC07756.1 | KEGG:R03876 | ubiquitin protein ligase, putative, |
| EEC20239.1 | KEGG:R03532 | thioredoxin-dependent peroxide reductase, which confer a protective role in cells through  its peroxidase activity by reducing hydrogen peroxide |
| EEC04717.1 | KEGG:R03876 | RNA polymerase II transcription elongation factor, putative, |
| EEC17038.1 | KEGG:R02541 | RAB-33, putative [Ixodes scapularis], exosome biogenesis and secretion between arthropods and mammals. |
| EEC09178.1 | KEGG:R07364 | acireductone dioxygenase, putative, Cysteine and methionine metabolism |
| EEC17038.1 | KEGG:R02541 | RAB-33, putative [Ixodes scapularis], exosome biogenesis and secretion between arthropods and mammals. |
| EEC08267.1 | KEGG:R00253 | glutamine synthetase, putative, |
| EEC09478.1 | KEGG:R02324 | nicotinamide riboside kinase, putative, Nicotinate and nicotinamide metabolism |
| EEC06118.1 | KEGG:R02530 | 4-hydroxyphenylpyruvate dioxygenase, putative, Pyruvate metabolism |
| EEC07756.1 | KEGG:R03876 | ubiquitin protein ligase, putative, |
| EEC07147.1 | KEGG:R05330 | short-chain dehydrogenase, putative, Primary bile acid biosynthesis, Biosynthesis of unsaturated fatty acids |
| EEC11438.1 | KEGG:R02541 | RAS-related protein, putative, exosome biogenesis and secretion between arthropods and mammals |
| EEC08160.1 | KEGG:R02541 | GTP-binding protein Rhes, putative, exosome biogenesis and secretion between arthropods and mammals. |
| EEC10220.1 | KEGG:R04188 | 4-aminobutyrate aminotransferase, Valine, leucine and isoleucine degradation |
| EEC07981.1 | KEGG:R01049 | ribose-phosphate pyrophosphokinase 1 putative, |
|  |  | rn00030 Pentose phosphate pathway rn00230 Purine metabolism rn01100 Metabolic pathways rn01110 Biosynthesis of secondary metabolites rn01120 Microbial metabolism in diverse environments rn01200 Carbon metabolism rn01230 Biosynthesis of amino acids |
| EEC08267.1 | KEGG:R00253 | glutamine synthetase putative, Arginine biosynthesis, Alanine, aspartate and glutamate metabolism, Glyoxylate and dicarboxylate, nitrogen metabolism, Microbial metabolism in diverse environments, Biosynthesis of amino acids |
| EEC05869.1 | KEGG:R02268 | cytochrome P450 putative, Arachidonic acid metabolism |
| EEC04717.1 | KEGG:R03876 | RNA polymerase II transcription elongation factor, putative |
| EEC08267.1 | KEGG:R00253 | glutamine synthetase putative, Arginine biosynthesis, Alanine, aspartate and glutamate metabolism, Glyoxylate and dicarboxylate, nitrogen metabolism, Microbial metabolism in diverse environments, Biosynthesis of amino acids |
| EEC07251.1 | KEGG:R02541 | RAS-related protein putative, exosome biogenesis and secretion between arthropods and mammals |
| EEC17749.1 | KEGG:R02541 | RAB-9 and, exosome biogenesis and secretion between arthropods and mammals |
| EEC09478.1 | KEGG:R02324 | nicotinamide riboside kinase (putative), Nicotinate and nicotinamide metabolism |
| EEC05808.1 | KEGG:R03876 |  |
| EEC02290.1 | KEGG:R00004 | secreted inorganic pyrophosphatase, putative, |
| EEC15523.1 | KEGG:R02265 | Microsomal prostaglandin E synthase 2, Arachidonic acid metabolism |
| EEC08267.1 | KEGG:R00253 | glutamine synthetase putative, Arginine biosynthesis, Alanine, aspartate and glutamate metabolism, Glyoxylate and dicarboxylate, nitrogen metabolism, Microbial metabolism in diverse environments, Biosynthesis of amino acids |
| EEC10220.1 | KEGG:R04188 | (S)-3-amino-2-methylpropionate transaminase, Valine, leucine and isoleucine degradation |
| EEC01042.1 | KEGG:R00310 | Ferrochelatase, Porphyrin and chlorophyll metabolism, Biosynthesis of secondary metabolites, Heme biosynthesis |

**Table S3. Gene-specific PCR and qRT-PCR primers used in this study.**

| **Gene** | **GenBank ID** | **Forward Primer (5'-3')** | **Reverse Primer (5'-3')** | **Size (bp)** |
| --- | --- | --- | --- | --- |
| *RPS4 (qPCR)* | DQ066214.1 | GGTGAAGAAGATTGTCAAGCAGAG | TGAAGCCAGCAGGGTAGTTTG | 80 |
| *flaB (qPCR)* | Stone et al., 2015 | GGG TCT CAA GCG TCT TGG | GAA CCG GTG CAG CCT GAG | 139 |
